# A rapid growth rate underpins the dominance of *Hanseniaspora uvarum* in spontaneous grape juice fermentations

**DOI:** 10.1101/2024.08.16.608365

**Authors:** Cristobal A. Onetto, Jane McCarthy, Simon A. Schmidt

## Abstract

*Hanseniaspora uvarum* is consistently observed as the dominant non-*Saccharomyces* species in spontaneous grape juice fermentations. However, the physiological mechanisms and physicochemical variables influencing the prevalence of *H. uvarum* over other non-*Saccharomyces* species remain unclear. We tested the physicochemical parameters contributing to *H. uvarum* dominance by inoculating a chemically diverse set of grape juices with a mock community whose composition was defined following a comprehensive microbial survey of spontaneous fermentations. Our findings indicated that the chemical composition of grape juice had minimal impact on the microbial dynamics of fermentation, with *H. uvarum* emerging as the dominant non-*Saccharomyces* species in nearly all conditions tested. Grape juice composition primarily influenced the total yeast abundance of the mock community. Flow cytometry analysis confirmed that *H. uvarum* has a faster growth rate than *Saccharomyces cerevisiae* and several other *Hanseniaspora spp*.. Moreover, its growth was not affected by the presence of *S. cerevisiae*, explaining its rapid dominance in spontaneous fermentations. The rapid growth of *H. uvarum* negatively impacted the growth of *S. cerevisiae*, with significant implications for fermentation performance and sugar consumption. The results of this study suggest that the fast growth rate of *H. uvarum* enables it to quickly dominate the grape juice environment during the early stages of fermentation. This physiological advantage indicates that the initial abundance of *H. uvarum* may be critical to the outcome of spontaneous fermentations, as evidenced by its direct impact on the growth of *S. cerevisiae* and fermentation performance.

## 1. Introduction

The surfaces of vines and grapes host naturally occurring, complex consortia of fungal and bacterial species that directly influence vine growth and overall health (Morrison-Whittle and Goddard, 2015; Müller et al., 2016; Taylor et al., 2014). These microbial communities can vary significantly in their composition, influenced by numerous factors such as grape variety, geographical location, climate, and viticultural practices (Bokulich et al., 2014; Gayevskiy and Goddard, 2012; Pinto et al., 2015).

During harvesting and processing, yeast are transferred into the grape must, where they play an important role in shaping fermentation dynamics and contributing to the sensory attributes of wine (Bagheri et al., 2018; Ciani et al., 2010). This influence is significant during uninoculated (’spontaneous’) fermentation, where the absence of commercial yeast strains allows microbial community members to impact the fermentation process directly. In spontaneous fermentations, a complex microbial succession of yeasts and yeast-like fungi is generally observed as the grape must ferments. Initially, a diverse mixture of aerobic and apiculate yeasts, which typically reside on the surface of intact grape berries, dominate the fresh grape must. However, most of these species rapidly decline due to the decreasing oxygen levels and increasing ethanol concentrations, allowing mildly fermentative species to increase in abundance (Beltran et al., 2002; Combina et al., 2005; Fleet, 1990, 2008; Fleet et al., 1984). The abundance and population dynamics of these species can be influenced by several abiotic parameters such as sulfur dioxide (SO_2_), temperature, pH, oxygen, and nutrient availability (Ciani et al., 2016; Cuijvers et al., 2020; Rollero et al., 2015; Varela et al., 2021).

Our recent microbial survey of more than 700 individual spontaneous fermentations revealed that once fermentation initiates, the complexity of these communities reduces considerably, with *H. uvarum* and *S. cerevisiae* emerging as the predominant yeast species during the initiation (1 Bé drop) and entire fermentation (10% sugar remaining) process in most of the ferments investigated (Onetto et al., 2024). This phenomenon has been consistently reported in earlier studies worldwide (Beltran et al., 2002; Ciani et al., 2016; Combina et al., 2005; Fleet, 1990). However, no studies specifically address the underlying mechanisms of *H. uvarum* dominance or the physicochemical variables that may influence the prevalence of this species over other non-Saccharomyces species.

Extensive research on the model yeast *S. cerevisiae* has established its high suitability for grape juice fermentation due to its superior fermentative ability, growth rate, and ethanol tolerance (Parapouli et al., 2020). In contrast, *H. uvarum* is a weak, Crabtree-negative fermentative species (Langenberg et al., 2017; Rodicio and Heinisch, 2017). From a genetic standpoint, *Hanseniaspora spp*., particularly *H. uvarum*, have a significantly reduced genome size compared to other budding yeasts in the subphylum Saccharomycotina and have lost several key cell-cycle and DNA repair genes (Steenwyk et al., 2019). Given these factors, further research on *H. uvarum* is required to fully uncover its competitive advantage during early fermentation, especially in comparison to other non-*Saccharomyces* yeast species found in grape must.

This study aimed to identify the parameters that modulate the microbial dynamics of spontaneous fermentation and investigate why *H. uvarum* consistently emerges as a dominant yeast under various conditions. We also aim to uncover the physiological mechanisms underlying the rapid abundance of this species in spontaneous fermentations.

## 2. Material and Methods

### 2.1. Mock community experiments

#### 2.1.1 Strains and mock community assembly

The strains used in this study are available in The Australian Wine Research Institute (AWRI) culture collection. Details on strain number and modifications are available in Table S1. To assemble the mock community, representative isolates for six species (Table S1) were selected based on their relative abundance at the initiation of fermentation reported in a recent survey of spontaneous fermentations in Australia (Onetto et al., 2024). Isolates were grown for 48 h in YPD (2 % glucose, 2 % peptone and 1 % yeast extract) and then counted using a Neubauer improved chamber (Weber Scientific, England). Each species was pooled into 50/50 grape juice:H_2_O to achieve a total inoculation rate of 10^4^ CFU/mL and an individual species inoculation rate as presented in Table S2, calculated based on the total average relative abundance for each species previously reported by Onetto et al. (2024) (Table S2).

#### 2.1.2 Grape juice and fermentation conditions

The growth dynamics of the mock community were assessed using a total of 8 different Chardonnay musts obtained from diverse grape-growing regions (Table S3). These juices were selected based on their chemical composition to cover a range of compositional parameters, i.e. yeast available nitrogen (YAN), pH, metal composition and sugar concentrations.

Determination of element concentrations for each juice by ICP-MS was performed by Affinity Labs (International Organization for Standardization 17025 accredited laboratory, Adelaide, SA, Australia). Samples were extracted in 2 mL HNO_3_ + 0.5 mL H_2_O_2_ for 90 min at 90 °C in screw-capped 50 mL polypropylene tubes (Environmental Express). Samples were diluted to a consistent volume of 20 mL with Milli-Q water and mixed thoroughly. Undissolved particles were allowed to settle, and then analysis was performed using a Perkin Elmer Nexion 350D ICP-MS with the KED mode, as previously described (Wheal and Wilkes, 2021). YAN was also determined by Affinity Labs as previously described by (Bergmeyer and Beutler, 1990; Bruce and Christian, 1998).

Each juice was sterile-filtered before being inoculated with the assembled mock community. All fermentations were performed in triplicate in 100 mL sealed Schott flasks fitted with gas release valves and stirred at 230 rpm at 18 °C. Samples were taken daily for DNA extraction and sugar (glucose + fructose) enzymatic determination as previously described (Hohorst, 1965) with adaptations as described by Vermeir et al. (2007). Total viable yeast cell concentrations were determined by dispensing 50 µL aliquots of serially diluted samples onto YPD (1 % w/v yeast extract, 2 % w/v peptone and 2 % w/v D-glucose) and Lysine agar plates using an automated spiral plater (WASP 2, Don Whitley Scientific, Australia). The agar plates were incubated at 27 °C for approximately 2–5 days and enumerated using a Protocol 3 colony counter (Synopsis, Don Whitley Scientific, Australia). Finished ferments were cold-settled for 7 days at 4 °C, and then supernatant samples were taken to quantify fermentation-derived volatile compounds as previously described (Onetto et al., 2020).

#### 2.1.2 ITS sequencing and data analysis

DNA was extracted from pellets using a Gentra puregene Yeast/Bact kit (Qiagen). PCRs were performed using 5 ng of DNA with primer sequences designed to amplify the fungal ITS region while adding experiment-specific inline barcodes. Amplification (30 cycles, 56°C annealing, 20 s extension, KAPA 2G Robust polymerase) targeting the ITS region was performed using primers BITS (ACCTGCGGARGGATCA) and B58S3 (GAGATCCRTTGYTRAAAGTT) (Bokulich and Mills, 2013). Samples were pooled and sequenced in a P2 Solo device (Oxford Nanopore Technologies, Oxford, UK) using the SQK-LSK114 kit. Pod5 files were base called using Guppy v. 6.5.7 (Oxford Nanopore Technologies, Oxford, UK) and then demultiplexed using cutadapt v4.4 (Martin, 2011). Reads were then mapped to an ITS reference set of the six yeast species using minimap2 v2.28 (Li, 2018) and coverage for each species was calculated using Bedtools v2.31.1 (Quinlan and Hall, 2010).

### 2.2. Dual-species experiments

Assessment of the growth rate for *H. uvarum* and *S. cerevisiae* in chemically defined grape juice media (CDM) (Schmidt et al., 2011) under single and co-inoculation was performed using an *S. cerevisiae* strain transformed with a blue fluorescent protein (BFP) gene to allow precise cell counting of each species by flow cytometry. Details on the transformation protocol are available in (Onetto et al., 2021). Each species was inoculated at a rate of 10^4^ cells/mL as determined by flow cytometry. Fermentations were carried out in 100 mL sealed Schott flasks fitted with gas release valves and stirred at 230 rpm at 18 °C. Cells were counted by flow cytometry using a Guava® easyCyte 12HT instrument (Merck Millipore, Burlington, MA, USA). Prior to cell counting, cells were diluted in PBS (NaCl 8 g/L, KCl 0.2 g/L, Na_2_HPO_4_ 1.44 g/L, KH_2_PO_4_ 0.24 g/L pH 7.4) to a concentration lower than 5 × 10^6^ cells/mL. Forward and side scatter detectors were used to determine particle size and estimate cell numbers. A minimum of 5000 events with a throughput lower than 500 event/μL was measured in all samples. A replicate experiment was carried out in parallel under the same conditions, using microplates to obtain absorbance measurements at 600 nm every 2 h.

## 3. Results and Discussion

### 3.1. Microbial dynamics of a mock-community in grape juice

Recent investigations into the fungal diversity on grapes (Liu and Howell, 2021) and the microbial dynamics of spontaneous grape juice fermentations (Bokulich et al., 2014; Onetto et al., 2024) have revealed a substantial diversity of fungal species inhabiting grape surfaces. Despite this microbial diversity, research has demonstrated that only a few species dominate during the initial stages of fermentation (Beltran et al., 2002; Ciani et al., 2016; Combina et al., 2005; Fleet, 1990; Onetto et al., 2024). Our recent investigation, encompassing more than 700 individual spontaneous fermentations, provided a much more detailed understanding of yeast diversity and relative abundance throughout the fermentation of grape must, and revealed that *H. uvarum* and *S. cerevisiae* are the predominant yeast species (Onetto et al., 2024). While *S. cerevisiae* is widely acknowledged as the principal species responsible for the completion and major contribution to the alcoholic fermentation process, high abundances of *H. uvarum* have been observed consistently throughout fermentations worldwide (Beltran et al., 2002; Combina et al., 2005; Heard and Fleet, 1985; Hierro et al., 2006; Holloway et al., 1990; Onetto et al., 2024; Snyder et al., 2024; Zott et al., 2010).

This study aimed to investigate the microbial dynamics of spontaneous fermentation in a simplified and controlled environment to ascertain whether *H. uvarum* would emerge as a dominant yeast species alongside *S. cerevisiae*. Utilising previously published relative abundance data (Onetto et al., 2024), we constructed a mock community to simulate the natural abundance of yeast species observed during the initiation of spontaneous fermentation. This community included *S. cerevisiae, H. uvarum*, and four other prevalent species: *Aureobasidium pullulans, Metschnikowia pulcherrima, Pichia kudriavzevii*, and *Torulaspora delbrueckii*. We tracked the microbial dynamics of these species throughout the fermentation of sterile Chardonnay grape must, employing a combination of plating and ITS-metabarcoding techniques. To stimulate a more competitive environment, we implemented a low total inoculation rate of 10^4^ CFU/mL, within the lower end of the expected yeast abundance of freshly pressed grape juice (Ribéreau-Gayon et al., 2006). Based on prior reports, we anticipated that *H. uvarum* and *S. cerevisiae* would dominate over other yeast species. Consequently, we included a treatment with only *H. uvarum* and *S. cerevisiae* at the same inoculation rate as the mock community to evaluate whether these species exhibited a different behaviour under a co-inoculation regime.

Given the known variations of ribosomal DNA (rDNA) locus copy number in yeast (Hotz et al., 2022; Mansisidor et al., 2018), we initially performed ITS sequencing on our assembled mock community to determine if discrepancies existed between the expected relative abundance, calculated based on cell counts, and the estimated relative abundance, determined through ITS metabarcoding. Although minor differences in the abundance of individual species were observed between the two methods, the six species maintained their rank in abundance using ITS-metabarcoding (Table S2), indicating that this methodology was sufficient for investigating the dynamics of these species throughout fermentation.

Despite its substantially lower inoculation rate compared to *S. cerevisiae*, ITS metabarcoding (Figure 1) and viable plating of the co-inoculation treatments at sugar consumption by one Be (T1; *S. cerevisiae*: 3.2×10^7^ - SD 1.9×10^7^, *H. uvarum*: 9.7×10^7^ – SD 1.0×10^7^) demonstrated that *H. uvarum* rapidly dominated under both mock and co-inoculation regimes and maintained a high abundance at 50% sugar consumption (T2; Figure 1). No differences in the rates of sugar consumption were observed between treatments, indicating that the presence of other species had minimal impact on the fermentation dynamics (Figure S1). Previous studies of inoculated (with *S. cerevisiae*) and uninoculated grape musts have identified *H. uvarum* (anamorph *Kloeckera apiculata*) as a dominant yeast not only at the initiation but also at later stages of fermentation (Beltran et al., 2002; Combina et al., 2005; Heard and Fleet, 1985; Hierro et al., 2006; Holloway et al., 1990; Onetto et al., 2024; Zott et al., 2010). The specific genetic factors contributing to the competitive fitness of *H. uvarum* in the grape juice environment remain unclear; however, recent investigations suggest that the loss of specific genes, including key cell-cycle regulators (Haase et al., 2024), may be an adaptive strategy enabling this species to undergo a more rapid cell cycle than *S. cerevisiae*, as previously reported in CDM (Charoenchai et al., 1998). While negative cell-to-cell contact-dependent interactions between *S. cerevisiae* and *H. uvarum* have been reported (Pietrafesa et al., 2020), our findings suggest these interactions are insufficient to inhibit the growth of *H. uvarum* at the inoculation rates used.

**Figure 1.**
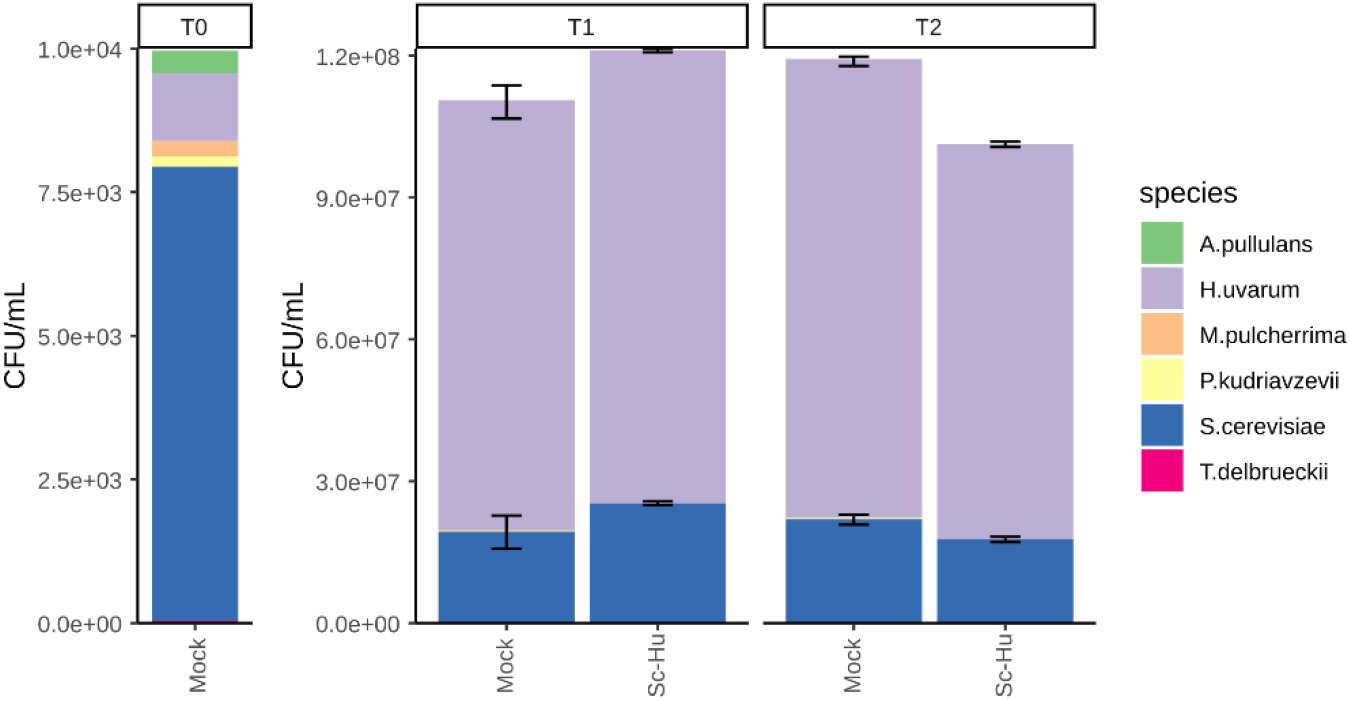
Microbial dynamics of a mock community and co-inoculation of *S. cerevisiae* and *H. uvarum* inoculated into sterile Chardonnay grape must. The relative abundance of each species was determined by ITS metabarcoding sequencing, and the total yeast abundance was determined by plating in YPD media. T0 represents the relative abundance of the mock community as determined by ITS sequencing. T1 and T2 represent the relative abundance of each species after 1 Be drop and 50% sugar consumption, respectively. Error bars show the standard deviation of the ITS relative abundance for the two most abundant species.

The chemical composition of grape juice has been shown to exert species-specific effects on the growth of wine yeast, influencing the microbial dynamics of mixed microbial communities (Ruiz et al., 2023). Consequently, in subsequent experiments, we evaluated the microbial dynamics of our mock community in a set of compositionally contrasting Chardonnay grape juices obtained from different wine regions.

### 3.2 The impact of grape juice composition on microbial dynamics

We analysed the chemical composition of 63 Chardonnay grape juices obtained from various winemaking regions and performed hierarchical clustering using the chemical compositional data (Figure S2). We selected eight juices from this set representing compositional differences observed in the major clusters (Figure S2, Table S3). The selected juices were inoculated with the mock community to assess whether grape juice composition impacts microbial dynamics throughout fermentation.

Assessment of total yeast growth at three time points and sugar consumption kinetics throughout fermentation revealed significant differences between juices (Figures 2 and 3). Fermentation duration (< 2 g/L of total sugars) varied substantially between treatments, ranging from 10 to 24 days. Furthermore, one juice exhibited minimal fermentation activity after 30 days of inoculation and contained high residual sugar (Figure 3).

**Figure 2.**
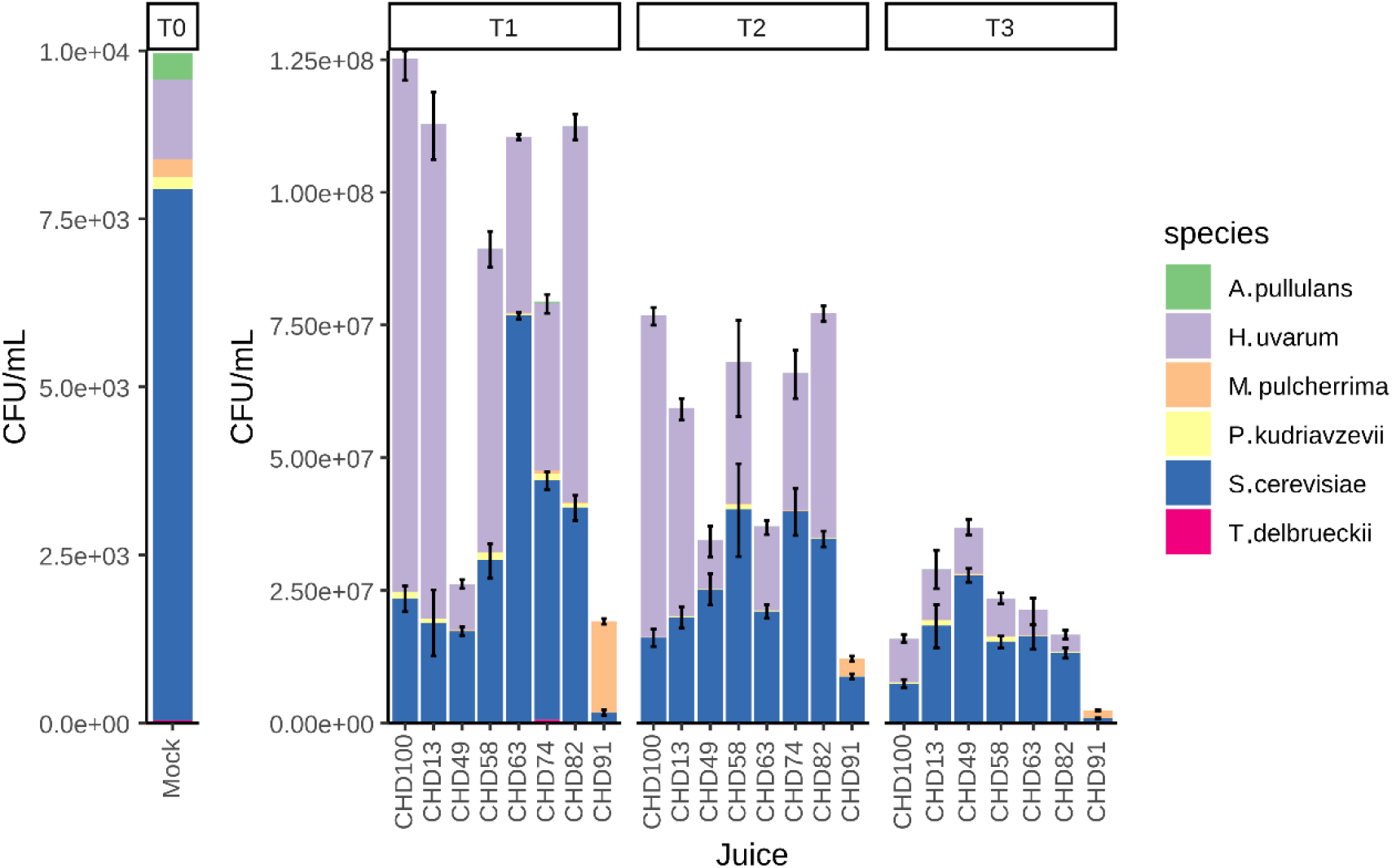
Microbial dynamics of a mock-community inoculated into eight sterile Chardonnay grape musts. The relative abundance of each species was determined by ITS metabarcoding sequencing, and the total yeast abundance was determined by plating. T0 represents the relative abundance of the mock-community as determined by ITS sequencing with a total inoculation rate of 10^4^ CFU/mL. T1, T2 and T3 represent the relative abundance of each species after 1 Be drop, 50% and 90% sugar consumption, respectively. Error bars show the standard deviation of the ITS relative abundance for the two most abundant species.

**Figure 3.**
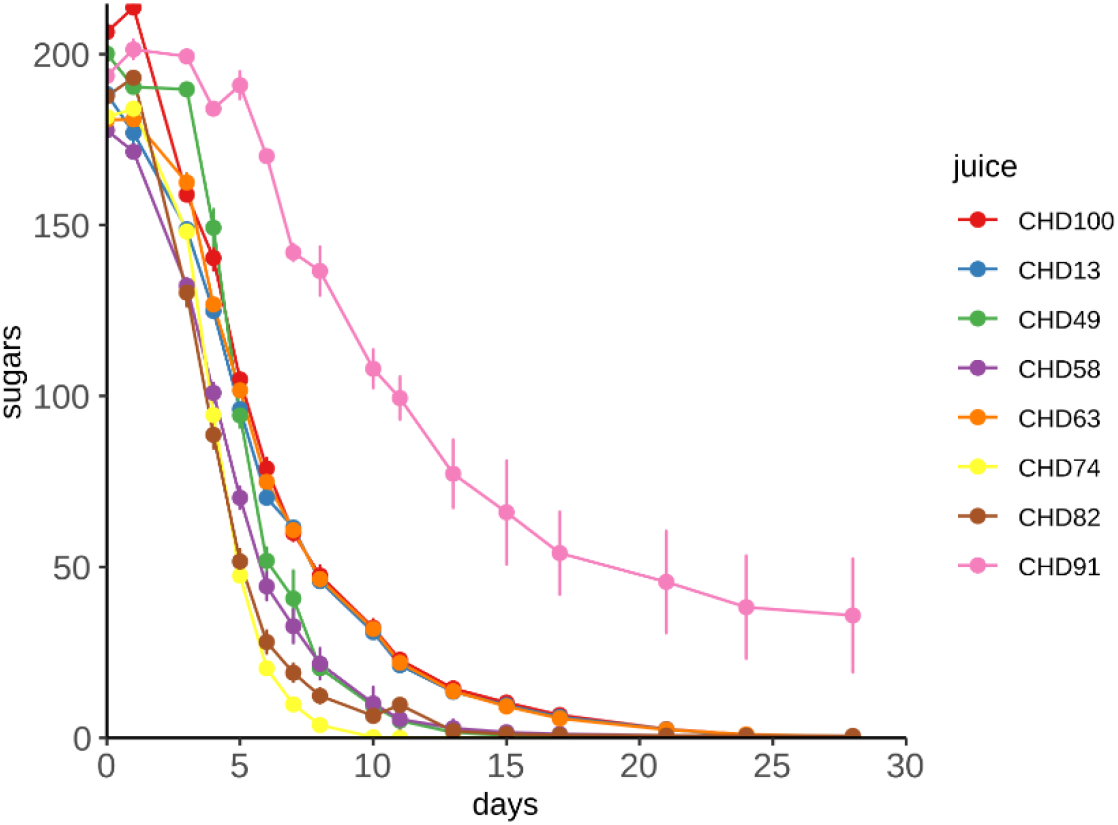
Sugar consumption of the mock community inoculated into eight different Chardonnay grape juices. Points show the mean of three replicates with error bars indicating standard deviation.

Consistent with the initial experiment, *H. uvarum* and *S. cerevisiae* were the two predominant species in all treatments except one, which was dominated by *M. pulcherrima* at T1 and showed the lowest total yeast growth and fermentation activity among all treatments (Figures 2 and 3). While the composition of each juice clearly influenced total yeast growth and fermentation kinetics, the individual microbial population dynamics of the mock community appeared consistent between juices, suggesting that the initial structure of the microbial population in spontaneous ferments is the primary determinant of microbial dynamics throughout fermentation. This observation aligns with the findings by Bagheri et al. (2020), who reported that absolute population numbers of each species are condition-dependent, but the broad trends in population dynamics are independent of the grape must chemical parameters tested. These results also further support our recent survey of 772 individual grape juices, where *S. cerevisiae* and *H. uvarum* were observed as the two dominant species throughout the entire fermentation in most of the juices investigated (Onetto et al., 2024). Despite this, *H. uvarum* is not always reported to dominate over *S. cerevisiae* in spontaneous fermentations. Our experiments represent a simplified model of what is expected to occur in natural spontaneous fermentation, where a higher diversity of species interact during the initial stages of fermentation. Increased species richness elevates the likelihood of observing negative interactions within the yeast community (Ruiz et al., 2023), which might explain the substantial variations we have previously observed in the abundance of *H. uvarum* at different fermentation stages (Onetto et al., 2024). These differences are likely attributable to a multifactorial interplay involving the chemical composition of the juice, the initial microbial abundance of each species, the complexity of the initial microbial community and the winemaking parameters used throughout the fermentation (e.g., temperature and SO_2_). Nevertheless, based on these results, it is evident that *H. uvarum* possesses a competitive advantage over other fungal species, enabling it to rapidly dominate the initial stages of fermentation in diverse conditions.

Extreme compositional parameters might still have a significant effect on particular species, as observed in juice CHD91, which had an unusually low pH of 2.9 and likely could not support successful spontaneous fermentation, allowing more resilient species such as *M. pulcherrima* (Al-Nijir et al., 2024; Santamauro et al., 2014) to survive. Furthermore, other winemaking-related variables, such as aeration and externally added SO_2,_ have been shown to induce shifts in microbial population dynamics throughout fermentation (Cuijvers et al., 2020; Varela et al., 2021).

Differences in the relative abundance of *S. cerevisiae* and *H. uvarum* were observed between juices, with four juices showing a higher abundance of *H. uvarum* at the initiation of fermentation. Of these, three juices (CHD100, CHD13, and CHD82) retained a higher abundance of *H. uvarum* at T2 (50% sugar consumption) and a single juice (CHD100) at 90% sugar consumption (T3; Figure 2). Consistent with several reports (Beltran et al., 2002; Combina et al., 2005; Onetto et al., 2024), *S. cerevisiae* was observed as the dominant species by the end of fermentation.

To understand the compositional characteristics impacting yeast growth, we performed a correlation analysis of the yeast growth data against each chemical parameter and the fermentation-derived volatiles of the resulting wines (Table S4). pH emerged as the strongest positively correlated parameter (r = 0.95, p < 0.001) with total yeast abundance, and was also correlated with the abundance of *H. uvarum, P. kudriavzevii*, and *T. delbrueckii* (Table S4); however, this parameter showed no correlation with fermentation duration, which was highly correlated (r = 0.91, p < 0.01) with the abundance of *S. cerevisiae*. The observed correlation between the time for fermentation completion and the abundance of *S. cerevisiae* is not surprising due to the strong fermentative ability of this organism and its known abundance in most spontaneously fermented wines. In contrast, *H. uvarum*, despite its high abundance, has poor fermentative capability (Langenberg et al., 2017) and is considered a Crabtree-negative yeast species (Rodicio and Heinisch, 2017). These results suggest that pH could be a critical parameter affecting the survival of these species in wine but might not impact the ability of *H. uvarum* to dominate in the early stages of fermentation within the pH ranges commonly observed in commercial wine production (Waterhouse et al., 2024).

As reported in several studies (Moreira et al., 2008; Plata et al., 2003), the abundance of *H. uvarum* was also correlated with the concentration of esters in the resulting wines, with ethyl acetate, ethyl butanoate, and 3-methylbutyl acetate concentrations showing a positive correlation with the abundance of *H. uvarum* at T1 and T2. It is for this particular reason that this species is mainly considered detrimental to wine production, especially when the concentrations of ethyl acetate exceed the range of 150-200 mg/L (Jackson, 2008). However, recent studies have shown large strain variability in the production of these volatiles, and some positive impacts on wine production have been reported (van Wyk et al., 2024).

Overall, these results indicate that the microbial dynamics of spontaneous fermentation are minimally affected by the chemical composition of the grape juice used in this study. Furthermore, these findings provide insight into the competitive advantage of *H. uvarum*, suggesting that this advantage is not related to any specific compositional parameter or nutritional component but may instead be attributed to a physiological advantage or an antagonistic mechanism.

### 3.3. A rapid growth rate allows *H. uvarum* to dominate the fermentation microbiome

Considering our initial results demonstrating that *H. uvarum* can rapidly dominate over other species in the mock-community under diverse conditions, we designed an experiment to investigate the growth rates of *H. uvarum* and *S. cerevisiae* in detail using flow cytometry. A genetically modified *S. cerevisiae* strain expressing BFP was used in co-inoculation with *H. uvarum*, allowing total and *S. cerevisiae* cell numbers to be determined. For this work, both species were inoculated at an initial concentration of 10^4^ cells/mL in single and co-inoculation formats. We then monitored their growth and fermentation kinetics at several time points during fermentation (Figure 4).

**Figure 4.**
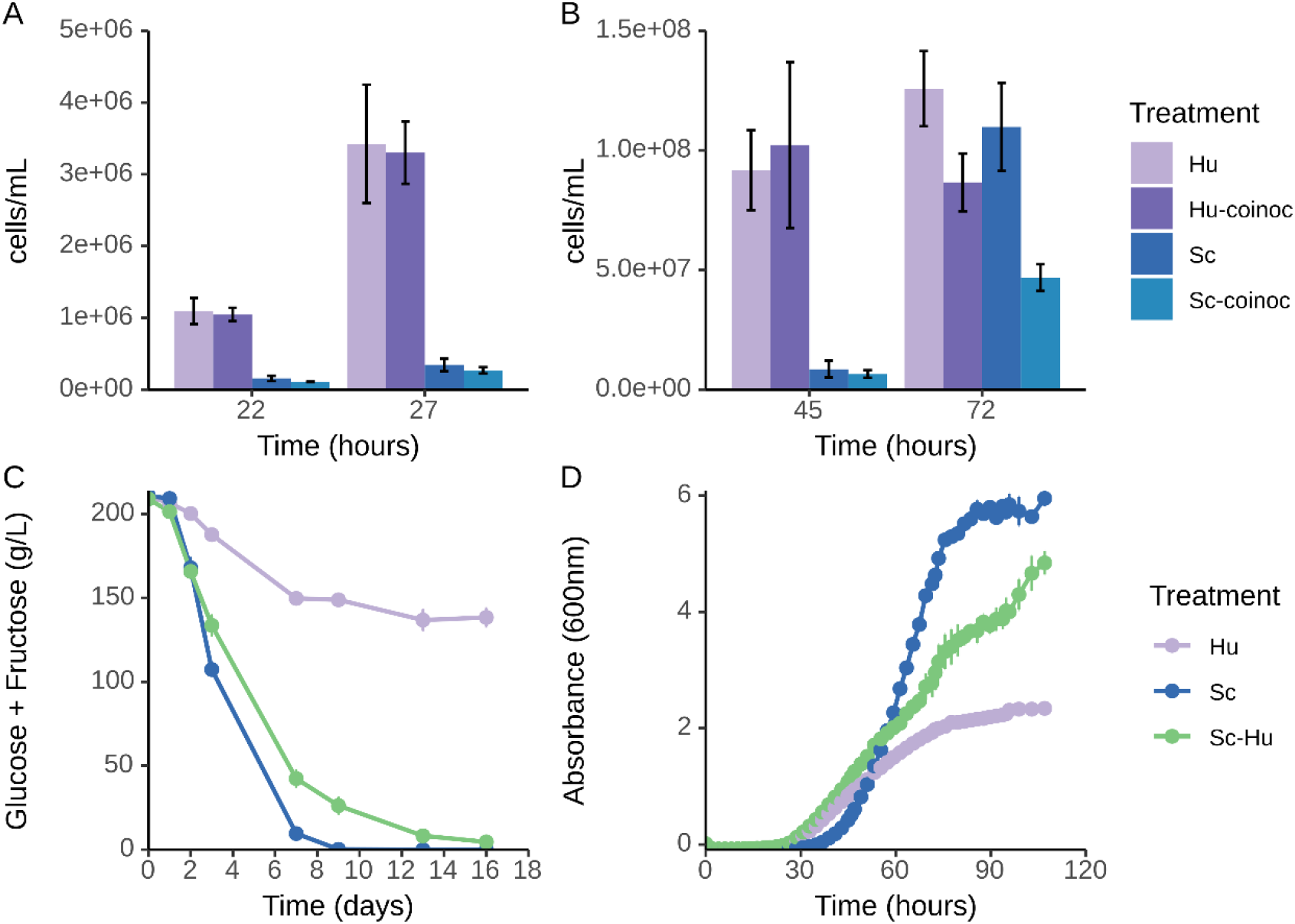
Growth kinetics of *H. uvarum* and *S. cerevisiae* in chemically defined grape juice media (CDM). Cell number determined by flow cytometry after 22 and 27 (A) and 45 and 72 hours (B) after inoculation of *H. uvarum* and *S. cerevisiae* at 10^4^ cells/mL into CDM. Coinoc treatments represent the cell number for *H. uvarum* (Hu) and *S. cerevisiae* (Sc) under co-inoculation. Sugar consumption (C) and growth kinetics (D) of the three treatments investigated. All data represent the mean of three replicates with error bars indicating standard deviation.

Flow cytometry cell counts at 22-, 27-, and 45-hours post-inoculation (Figure 4A and B) revealed significant differences between species (Table S5), with *H. uvarum* displaying a 7-, 10-, and 11-fold higher cell number compared to *S. cerevisiae*, respectively. No significant differences were observed for each species under single and co-inoculation regimes at these time points, indicating the absence of negative interactions between these two species that might impact growth. However, at 72 hours, a significant impact on the growth of *S. cerevisiae* under co-inoculation was observed (Table S5; Figure 4B and D), with clear implications for fermentation performance, sugar consumption, and fermentation completion in the co-inoculated treatments (Figure 4C). At this time point, *S. cerevisiae* had already reached a cell number similar to that of *H. uvarum* in the single species treatments (Figure 4B).

Log-transformed cell numbers obtained during the first three time points indicated that both species were in the exponential growth phase. Therefore, we used these data points to calculate the growth rates of these species. In agreement with earlier reports (Charoenchai et al., 1998), *H. uvarum* exhibited a faster growth rate (0.25 h^-1^, doubling time 2.7 hours) compared to *S. cerevisiae* (0.17 h^-1^, doubling time 3.9 hours), and was only minimally affected by the presence of *S. cerevisiae* in co-inoculation (Figure 4B). These results help to explain why *H. uvarum* is so prevalent throughout fermentation. Its fast growth rate allows this species to rapidly dominate the grape juice environment during the early stages of fermentation. Due to this physiological advantage, the initial abundance of *H. uvarum* might be critical to the outcome of spontaneous fermentations, as evidenced by the direct impact on the growth of *S. cerevisiae*. Previous investigations have shown that *H. uvarum* does not have high nitrogen requirements, with high concentrations of YAN still bioavailable even at the end of fermentation in both single and co-inoculation with *S. cerevisiae* (Ciani et al., 2006; Roca-Mesa et al., 2020). This suggests that nitrogen competition is not necessarily the reason for the observed impact on the growth of *S. cerevisiae*, and it might instead be attributed to the depletion of other nutrients such as thiamine (Bataillon et al., 1996), particularly due to the inability of *Hanseniaspora spp*. to synthesise this vitamin (Steenwyk et al., 2019). Furthermore, our results indicate that *S. cerevisiae* is unlikely to affect the growth of *H. uvarum* during the initial stages of spontaneous fermentation, in contrast to previously reported under co-inoculation (Pietrafesa et al., 2020).

Several *Hanseniaspora spp*. are commonly found on the surface of grapes, however, during fermentation, they are not as abundant as *H. uvarum*. To investigate whether other *Hanseniaspora spp*. possess a similar physiological advantage, we determined the growth rate in CDM at 28°C of *Hanseniaspora opuntiae, Hanseniaspora pseudoguilliermondii, Hanseniaspora guilliermondii*, and *Hanseniaspora valbyensis*, all belonging to the previously defined fast-evolving lineage (FEL) (Steenwyk et al., 2019). Among these species, only *H. guilliermondii* (0.51 h^-1^) exhibited a growth rate comparable to *H. uvarum* (0.425 h^-1^) (Table S6). Despite this, *H. guilliermondii* is rarely observed as a dominant species in spontaneously fermented wines. A possible explanation might be that *H. guilliermondii* is initially present at very lower numbers on the surface of grapes and therefore cannot compete with the initially high abundance of *H. uvarum* present on grapes (Liu and Howell, 2021) that is subsequently transferred into the grape juice (Onetto et al., 2024).

## 4. Conclusions

While a large diversity of yeast species is present in freshly pressed grape juice, research has demonstrated that, over time, only a few key species rapidly dominate the fermentation environment. One of these key species is *H. uvarum*, which is consistently reported as the predominant non-Saccharomyces species in spontaneous grape juice fermentations. This study aimed to identify the parameters that modulate the microbial dynamics in spontaneous fermentation and investigate whether *H. uvarum* would emerge as a dominant yeast under various conditions. Using a mock community based on a comprehensive microbial survey of spontaneous fermentations, we tested this hypothesis by inoculating a chemically diverse set of grape juices. *H. uvarum* was observed as the dominant non-Saccharomyces species in nearly all the conditions tested, maintaining high abundance throughout fermentation. This suggests that initial microbial community composition is the primary determinant of microbial dynamics during fermentation. In contrast, the chemical parameters of the grape must, primarily influences total yeast abundance.

Using flow cytometry, we confirmed that *H. uvarum* has a fast growth rate and is not significantly affected by the presence of *S. cerevisiae*, explaining its rapid dominance in spontaneous fermentations. Furthermore, *H. uvarum* had a faster growth rate than several other *Hanseniaspora spp*. commonly isolated from grapes.

From an adaptive viewpoint, *H. uvarum* appears less genetically equipped than other *Hanseniaspora spp*. for fermentation due to a notable loss of genes related to the cell cycle and DNA repair (Steenwyk et al., 2019). Therefore, future research should focus on identifying the key genetic factors that enable *H. uvarum* to achieve a rapid growth rate and competitive advantage over other non-Saccharomyces yeast species found in grape must.

## Acknowledgements

This work was supported by Wine Australia, with levies from Australia’s grape growers and winemakers and matching funds from the Australian Government. The AWRI is a member of the Wine Innovation Cluster (WIC) in Adelaide.

## 6. Supplementary Figures

**Figure S1.**
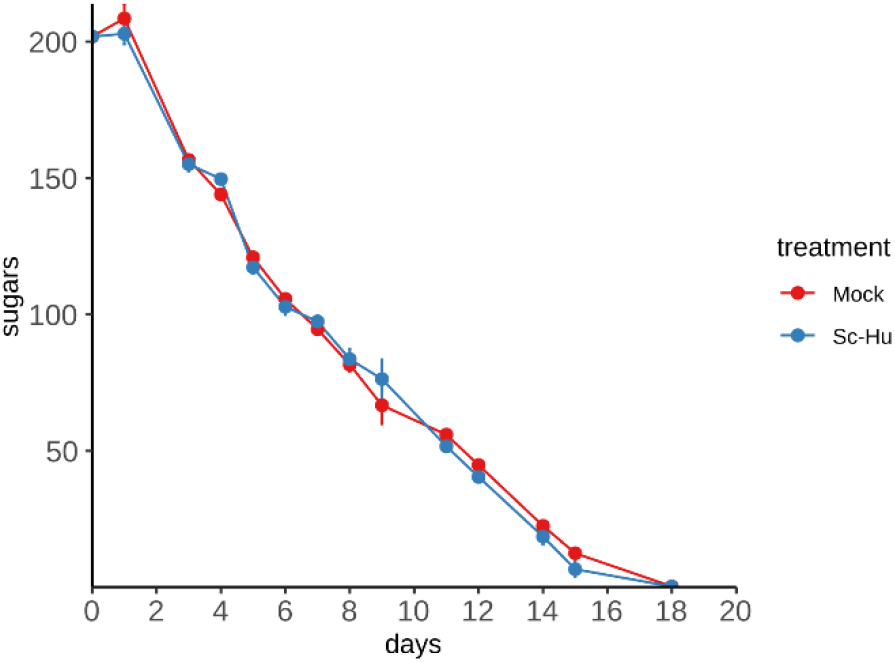
Sugar consumption of the mock community and *S. cerevisiae*-*H. uvarum* co-inoculated into a Chardonnay grape juice. Points show mean of three replicates with error bars indicating standard deviation.

**Figure S2.**
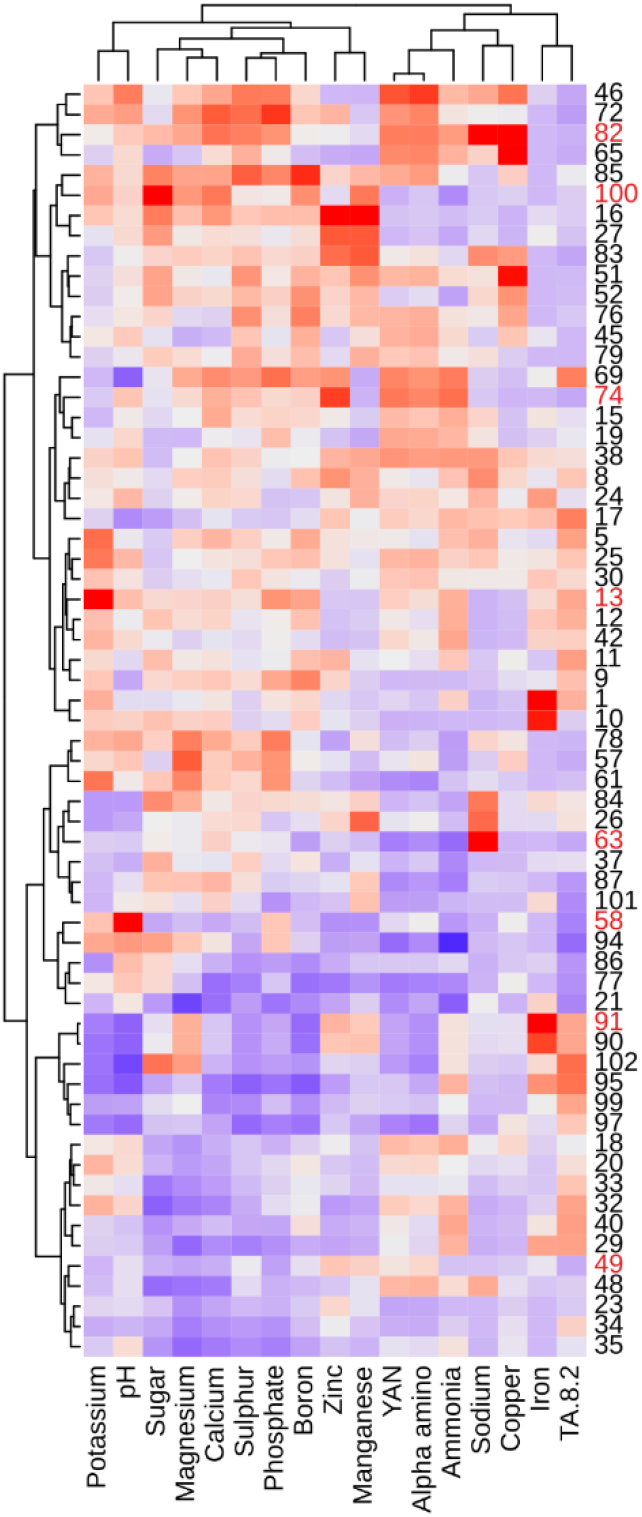
Heatmap displaying the hierarchical clustering of 63 Chardonnay grape juices based on their chemical composition. The row numbers in red represent the eight juices (Table S3) selected for fermentation.

